# Multimodal representations of person identity individuated with fMRI

**DOI:** 10.1101/090472

**Authors:** Stefano Anzellotti, Alfonso Caramazza

## Abstract

Recognizing the identity of a person is fundamental to guide social interactions. We can recognize the identity of a person looking at her face, but also listening to her voice. An important question concerns how visual and auditory information come together, enabling us to recognize identity independently of the modality of the stimulus. This study reports converging evidence across univariate contrasts and multivariate classification showing that the posterior superior temporal sulcus (pSTS), previously known to encode polymodal visual and auditory representations, encodes information about person identity with invariance within and across modality. In particular, pSTS shows selectivity for faces, selectivity for voices, classification of face identity across image transformations within the visual modality, and classification of person identity across modality.

## Introduction

Recognizing a person's identity is rapid and effortless, but the neural mechanisms that enable us to do it are complex. Two crucial sources of information we rely on to recognize others are the visual appearance of the face and the sound of the voice. What are the neural mechanisms that enable us to recognize the identity of a person across both face and voice stimuli?

Neuropsychological and neuroimaging studies implicated regions in occipitotemporal cortex and in the anterior temporal lobes (ATL) in face recognition (Meadows 1974; Tranel et al. 1997; Sergent et al. 1992; Kanwisher et al. 1997; Rajimehr et al. 2008, Thomas et al. 2009 for a review see Collins and Olson 2014). Recent research shows that patterns of response in these regions distinguish between images of faces (“face tokens”) that depict different people (Kriegeskorte et al. 2007; Nestor et al. 2011; Anzellotti et al. 2013; Goesaert et al. 2013; Verosky et al. 2014; Cowen et al. 2014; Natu et al. 2010; Anzellotti and Caramazza 2015; for a review see Anzellotti and Caramazza 2014), and do so with some invariance across changes in facial expression (Nestor et al. 2011), viewpoint (Anzellotti et al. 2013) and face parts (Anzellotti et al. 2015).

In parallel with this research, neuroimaging experiments found stronger responses to voices than to speech envelope noises in anterior portions of the superior temporal gyrus (STG) and in a more posterior region at the juncture between the superior temporal sulcus/gyrus and the parietal lobe (von Kriegstein et al. 2003). Anterior and posterior STG also show stronger responses when participants attend to the identity of the speaker than to the speech content (von Kriegstein et al. 2003), and patterns of response in these regions encode information about the identity of a speaker (Formisano et al. 2008). In addition, recent studies (Watson et al. 2014a,b) found responses to faces and voices in the pSTS.

However, these studies leave several questions unanswered. In particular, the studies (Watson et al. 2014a,b) do not test whether pSTS encodes information that is sufficiently specific to distinguish between individual faces. Given the involvement of neighboring pSTS regions in emotion processing, for instance (Peelen et al. 2010, Skerry et al. 2014), the effects reported (Watson et al. 2014a,b) might derive from representations of information about faces and voices that is orthogonal to identity, for instance emotion information (see Peelen et al. 2010, Skerry and Saxe 2014). Second, one of the studies (Watson et al. 2014a) only analyzed the overall response to faces and voices, therefore it remains unclear whether face and voice stimuli elicit similar patterns of response, or whether they activate very different representations with different spatial patterns generated by different neural populations that coexist within a same region. The other study attempted to overcome this limitation using adaptation (Watson et al. 2014b), but adaptation across different stimuli can occur in the absence of representations that generalize across those stimuli, for instance as a consequence of attentional effects (see Anzellotti et al. 2014 for a more detailed discussion). As an extreme example, adaptation for the same face across viewpoints has been observed in early visual cortex (Mur et al. 2010), which is far from encoding view-invariant representations of faces.

Where is visual and auditory information integrated into a multimodal representation of person identity with sufficient specificity to distinguish between individuals and sufficient invariance to generalize across faces and voices? A candidate area for the convergence of face and voice identity is the anterior temporal lobe (ATL). Damage to ATL can lead to multimodal deficits (Hodges et al. 1992, Bozeat et al. 2000, Rogers et al. 2004, Coccia et al. 2004), including deficits for person recognition and naming (Damasio et al. 1996, Tranel et al. 1997). Furthermore, ATL contains both ventral face-selective (Rajimeher et al. 2008) and more dorsal (anterior STG) voice selective regions (von Kriegstein et al. 2003), although it remains unclear whether or not they overlap. Another candidate area of convergence is the posterior superior temporal sulcus (pSTS), previously implicated in visuo-auditory integration (Benevento et al. 1977, Hikosaka et al. 1988, Beauchamp et al. 2004). Posterior STS contains a mosaic of visually and auditorily responsive patches (Beauchamp et al. 2004), and many pSTS neurons respond to both visual and auditory stimuli (Benevento et al. 1977, Hikosaka et al. 1988). Portions of pSTS are face-selective (Grill-Spector et al. 1999) and are located near voice-selective regions in pSTG (von Kriegstein et al. 2003). However, it remains to be clarified whether the face-selective pSTS also shows voice-selective responses. Most importantly, it is unknown whether ATL and pSTS encode representations of person identity that generalize across different modalities, that could be used to support multimodal person recognition.

The present study investigated the representation of person identity in ATL and pSTS, presenting both face and voice stimuli within the same participants. The study comprises a univariate investigation of voice selectivity in face selective regions and of face selectivity in voice selective regions, and a multivariate analysis within face- and voice-selective regions of information about person identity that generalizes across modality. In the multivariate analysis, a classifier was trained to distinguish between identities in one modality and tested on its accuracy in distinguishing between those same identities in the other modality. The face-selective right pSTS was found to respond differentially to both faces over houses, and voices over tool sounds, and to encode information about person identity generalizing across modality. If the right pSTS encodes information of person identity, it should classify identity also generalizing across transformations in the visual modality, such as changes in viewpoint. A reanalysis of an earlier study, in which the right pSTS had not been investigated (Anzellotti et al. 2013), revealed that patterns of response in the right pSTS could indeed classify between different identities generalizing across viewpoints. These results indicate that the right pSTS encodes information about person identity generalizing both within and across different modalities.

## Materials and Methods

### Participants and stimuli

Eleven volunteers (all Italian native speakers; 6 female; age range: 19-32, mean = 24) took part in the experiment. Three famous Italian politicians (Matteo Renzi, Pierluigi Bersani, and Silvio Berlusconi) were chosen to generate the stimuli. Two grayscale images of each face were selected and cropped to an oval (see Fig. 1), and equated in luminance and contrast. Two audio clips were selected for each of the three politicians: one in which they said “Italia” (“Italy”) and one in which they said “governo” (“government”). The audio stimuli were further equated in loudness.

**Figure 1.**
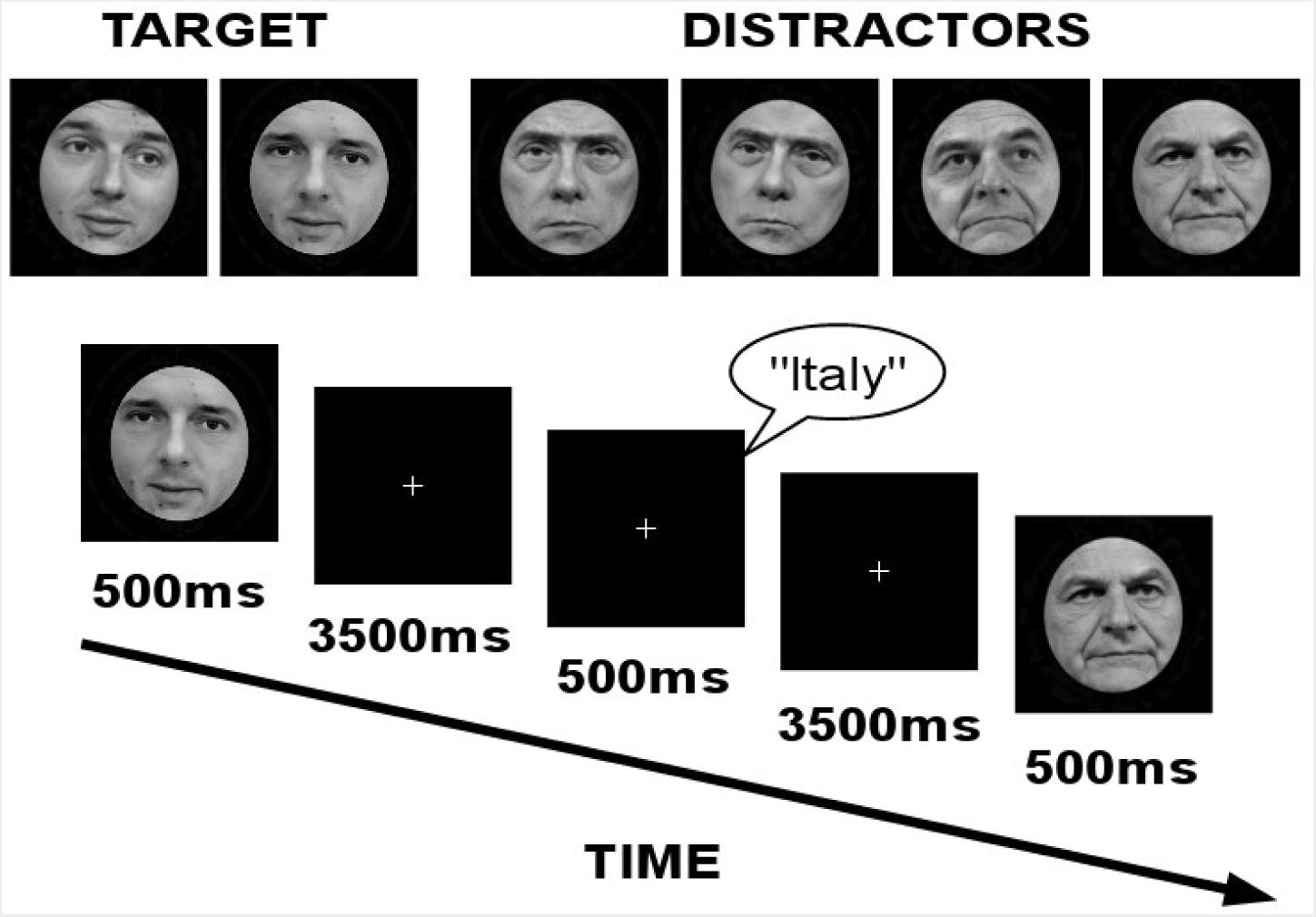
Face stimuli used in the experiment and experimental paradigm. Three famous politicians were selected, and two different pictures of each individual were presented. Stimuli were presented for 500ms followed by a 3500ms fixation.

### Behavioral training

On the day before the fMRI experiment, participants took part in a training session in which they watched an interview (lasting approximately 25 minutes) with each of the three individuals. After watching the video, they completed a 40-minute behavioral task in which they saw the six face images and heard the six voice sounds and could practice discriminating between the different identities.

### Experiment design

Inside the scanner, participants completed two localizer runs (a face localizer and a voice localizer) and five experimental runs. Before entering the scanner, participants were instructed to consider a given individual (e.g., Matteo Renzi) as the target. Participants were instructed to press a button with the index finger of the right hand when the target was presented, and a button with the middle finger of the right hand when a distractor was presented, irrespectively of stimulus modality. In the face localizer, participants were shown 16 seconds blocks of faces and houses, and performed a 1-back task reporting whether a stimulus was identical to the one that had been presented in the previous trial. In the voice localizer, participants heard 16 seconds blocks of voices and tool sounds, and performed an analogous 1-back task. In each experimental run, each “distractor” face and each “distractor” voice was presented 16 times, while the target face and target voice were presented for 8 trials. Given that there are two “distractor” identities and one “target” identity, this implies that the target identity was presented on 20% of the trials. Stimuli were presented for 500ms and were followed by a 3500ms fixation. The order of the trials was optimized to maximize efficiency using Optseq 2 (http://surfer.nmr.mgh.harvard.edu/optseq/).

### MRI data acquisition

MRI data were collected on a Bruker BioSpin MedSpec 4T at the Center for Mind/Brain Sciences (CIMeC) of the University of Trento using a USA Instruments eight-channel phased-array head coil. Before collecting functional data, a high-resolution (1×1×1 mm3) T1-weighted MPRAGE sequence was performed (sagittal slice orientation, centric phase encoding, image matrix = 256 × 224 [Read × Phase], field of view =256 × 224 mm [Read × Phase], 176 partitions with 1-mm thickness, GRAPPA acquisition with acceleration factor = 2, duration = 5.36 minutes, repetition time = 2700, echo time = 4.18, TI = 1020 msec, 7° flip angle).

Functional data were collected using an echo-planar 2D imaging sequence with phase oversampling (image matrix = 70 × 64, repetition time = 2000 msec, echo time = 21 msec, flip angle = 76°, slice thickness = 2 mm, gap = 0.30 mm, with 3 × 3 mm in plane resolution). Over four runs, 1260 volumes of 43 slices were acquired in the axial plane aligned along the long axis of the temporal lobe. Each run was preceded by a point-spread function (PSF) sequence, and distortion correction was implemented with the method described by Zaitsev and colleagues (2004).

### Data analysis

Data were preprocessed with SPM8 (http://www.fil.ion.ucl.ac.uk/spm/software/spm8/) running on MATLAB 2011a, and with custom MATLAB software using the MATLAB bioinformatics toolbox and LIBSVM (Chang and Lin 2011). Results were displayed with MRIcron (Rorden and Brett 2000).

The first 4 volumes of each run were discarded and slice-acquisition delays were corrected using the middle slice as reference. All images were corrected for head movement and normalized to the standard SPM8 EPI template. The BOLD signal was high-pass filtered at 128s and prewhitened using an autoregressive model AR(1).

Localizer data were smoothed with an isotropic 6mm full width at half maximum (FWHM) Gaussian kernel and modeled with a standard GLM. Regressors for the stimulus categories (faces and houses in the first localizer, human voices and tool sounds in the second localizer) were convolved with the SPM default haemodynamic response function (HRF). In addition to regressors for the stimulus categories, the model included six motion regressors and a regressor for the mean of the run. The contrast between faces and houses, and the contrast between human voices and tool sounds, were used to individuate peaks of selectivity in each individual participant. Functional ROIs were defined as 9mm radius spheres centered in the peaks of selectivity (see Tables 1 and 2 for the MNI coordinates of the peaks in each individual subjects for the face and voice localizers respectively). The ROIs defined with one localizer were used to investigate the difference between the responses to the two categories in the other localizer using a t-test on the differences between the mean ROI betas for the two categories across participants. The grey matter mask used for RFE was generated averaging the segmented grey matter masks from an independent group of participants (Anzellotti et al. 2015), smoothing the average grey matter, and setting a threshold to generate 0/1 map.

**Table 1.**
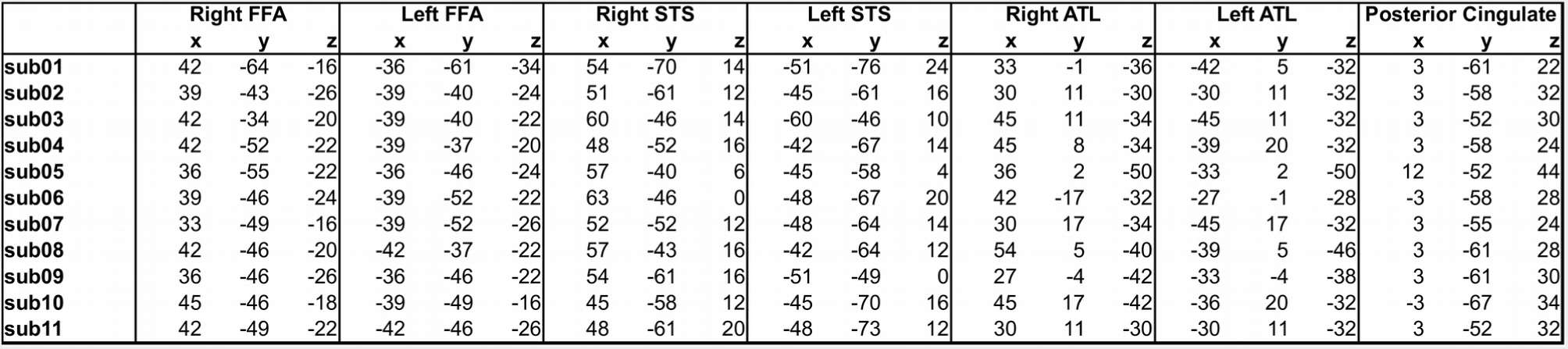
MNI coordinates of the centers of the spherical ROIs individuated with the faces>houses contrast.

|  | Right FFA |  |  | Left FFA |  |  | Right STS |  |  | Left STS |  |  | Right ATL |  |  | Left ATL |  |  | Posterior Cingulate |  |  |
| --- | --- | --- | --- | --- | --- | --- | --- | --- | --- | --- | --- | --- | --- | --- | --- | --- | --- | --- | --- | --- | --- |
|  | x | y | z | x | y | z | x | y | z | x | y | z | x | y | z | x | y | z | x | y | z |
| sub01 | 42 | -64 | -16 | -36 | -61 | -34 | 54 | -70 | 14 | -51 | -76 | 24 | 33 | -1 | -36 | -42 | 5 | -32 | 3 | -61 | 22 |
| sub02 | 39 | -43 | -26 | -39 | -40 | -24 | 51 | -61 | 12 | -45 | -61 | 16 | 30 | 11 | -30 | -30 | 11 | -32 | 3 | -58 | 32 |
| sub03 | 42 | -34 | -20 | -39 | -40 | -22 | 60 | -46 | 14 | -60 | -46 | 10 | 45 | 11 | -34 | -45 | 11 | -32 | 3 | -52 | 30 |
| sub04 | 42 | -52 | -22 | -39 | -37 | -20 | 48 | -52 | 16 | -42 | -67 | 14 | 45 | 8 | -34 | -39 | 20 | -32 | 3 | -58 | 24 |
| sub05 | 36 | -55 | -22 | -36 | -46 | -24 | 57 | -40 | 6 | -45 | -58 | 4 | 36 | 2 | -50 | -33 | 2 | -50 | 12 | -52 | 44 |
| sub06 | 39 | -46 | -24 | -39 | -52 | -22 | 63 | -46 | 0 | -48 | -67 | 20 | 42 | -17 | -32 | -27 | -1 | -28 | -3 | -58 | 28 |
| sub07 | 33 | -49 | -16 | -39 | -52 | -26 | 52 | -52 | 12 | -48 | -64 | 14 | 30 | 17 | -34 | -45 | 17 | -32 | 3 | -55 | 24 |
| sub08 | 42 | -46 | -20 | -42 | -37 | -22 | 57 | -43 | 16 | -42 | -64 | 12 | 54 | 5 | -40 | -39 | 5 | -46 | 3 | -61 | 28 |
| sub09 | 36 | -46 | -26 | -36 | -46 | -22 | 54 | -61 | 16 | -51 | -49 | 0 | 27 | -4 | -42 | -33 | -4 | -38 | 3 | -61 | 30 |
| sub10 | 45 | -46 | -18 | -39 | -49 | -16 | 45 | -58 | 12 | -45 | -70 | 16 | 45 | 17 | -42 | -36 | 20 | -32 | -3 | -67 | 34 |
| sub11 | 42 | -49 | -22 | -42 | -46 | -26 | 48 | -61 | 20 | -48 | -73 | 12 | 30 | 11 | -30 | -30 | 11 | -32 | 3 | -52 | 32 |

**Table 2.** MNI coordinates of the centers of the spherical ROIs individuated with the voices>tool sounds contrast.

|  | DMPFC |  |  | Right pSTS |  |  | Left pSTS |  |  | Right midSTS |  |  | Left midSTS |  |  | Right aSTS |  |  | Left aSTS |  |  | Posterior Cingulate |  |  |
| --- | --- | --- | --- | --- | --- | --- | --- | --- | --- | --- | --- | --- | --- | --- | --- | --- | --- | --- | --- | --- | --- | --- | --- | --- |
|  | x | y | z | x | y | z | x | y | z | x | y | z | x | y | z | x | y | z | x | y | z | x | y | z |
| sub01 | 0 | 62 | -12 | 54 | -67 | 12 | -45 | -70 | 28 | 66 | -19 | 0 | -63 | -25 | 6 | 57 | 11 | -14 | -51 | 11 | -14 | -6 | -61 | 22 |
| sub03 | 9 | 59 | -26 | 54 | -70 | 32 | -39 | -55 | 16 | 63 | -16 | -4 | -66 | -13 | -6 | 57 | 11 | -12 | -57 | 11 | -16 | 0 | -70 | 32 |
| sub04 | 3 | 62 | -8 | 57 | -64 | 22 | -36 | -58 | 26 | 63 | -25 | -2 | -66 | -22 | 0 | 57 | 11 | -16 | -57 | 5 | -12 | 0 | -58 | 32 |
| sub05 | -3 | 53 | -6 | 51 | -67 | 20 | -39 | -70 | 18 | 57 | -16 | -4 | -57 | -40 | 6 | 57 | 8 | -14 | -57 | 2 | -12 | -6 | -61 | 28 |
| sub08 | 6 | 50 | -16 | 54 | -61 | 30 | -54 | -52 | 22 | 48 | -34 | 0 | -66 | -28 | -2 | 51 | 23 | -20 | -57 | 11 | -12 | -6 | -61 | 36 |
| sub12 | 6 | 50 | -20 | 39 | -58 | 22 | -54 | -55 | 12 | 63 | -10 | -12 | -63 | -16 | -10 | 57 | 8 | -18 | -54 | 5 | -16 | 0 | -58 | 26 |
| sub13 | 3 | 47 | -26 | 39 | -67 | 24 | -36 | -61 | 24 | 60 | -13 | -8 | -66 | -37 | 6 | 54 | 11 | -20 | -54 | 8 | -18 | -3 | -55 | 26 |
| sub14 | 6 | 53 | -12 | 45 | -73 | 32 | -51 | -67 | 18 | 63 | -13 | -4 | -63 | -13 | 4 | 57 | 8 | -16 | -54 | 2 | -10 | -9 | -61 | 38 |
| sub15 | 0 | 59 | -8 | 54 | -70 | 26 | -57 | -61 | 6 | 66 | -7 | -8 | -66 | -22 | -4 | 54 | 14 | -16 | -54 | 8 | -22 | 12 | -70 | 20 |
| sub16 | -6 | 56 | -12 | 45 | -55 | 14 | -48 | -67 | 14 | 63 | -10 | -8 | -60 | -19 | 0 | 57 | 8 | -12 | -60 | 2 | -10 | -6 | -67 | 32 |
| sub17 | 6 | 62 | -8 | 57 | -67 | 22 | -51 | -67 | 22 | 66 | -31 | 0 | -66 | -31 | 2 | 57 | 8 | -14 | -54 | 2 | -10 | 3 | -58 | 28 |

Experimental runs were modeled with a standard GLM that included separate regressors for each stimulus (convolved with SPM's HRF) as well as six motion regressors and a regressor for the mean of each run. Regions encoding information that distinguishes between face tokens depicting different people and regions encoding information that distinguishes between voice tokens corresponding to different people were identified with recursive feature elimination mapping (RFE mapping, Formisano et al. 2008; DeMartino et al. 2008). Recursive feature elimination mapping individuates the voxels that contribute most to a given classification using the weights of support vector machines (SVMs). More specifically, a support vector machine (SVM) is trained to perform the classification between two stimuli using the data from all voxels in the gray matter in one experimental session. Each voxel is assigned a weight, which reflects the extent to which it contributes to the classification. The voxels with the smallest weights are removed, and the procedure is repeated in training a new SVM to perform the classification using only the remaining voxels. This procedure is iterated until a given number of voxels is reached. The number of voxels was set to approximately half of the total number of voxels in the mask (in this case gray matter) so as to maximize the number of possible combinations of voxel locations (given by the binomial coefficient). Selected voxels are assigned a value of 1, while non-selected voxels are assigned a value of 0. Minimal gaussian smoothing is applied (3mm FWHM) in order to account for small differences across sessions and participants. The resulting maps for the different sessions and participants are averaged. This procedure yields an average "frequency map" with values between 0 and 1 for each voxel. The value for a particular voxel indicates how frequently that voxel is individuated as being among the most informative for the particular classification tested, which indicates how reliably that voxel contributes to the classification.

Significance is tested generating statistical thresholds with Monte Carlo simulations. Simulated data of the same dimension as the experimental data are generated for each session and participant. To account for spatial autocorrelation in the real data, the simulated data are smoothed with a gaussian kernel of FWHM identical to the smoothness of the real data measured with SPM. Then, the simulated data are processed with the same RFE mapping algorithm applied to the real data. The same mild smoothing (3mm FWHM) is applied to the maps obtained from the simulated data. The maps for different sessions and participants are averaged, using an identical procedure as that used for the real data, yielding a "frequency map" as in the case of the real data. This procedure is iterated for 200 simulations of the data. For each of the 200 frequency maps obtained, the maximum value in the map is stored in a 200-dimensional vector. Voxels in the frequency map obtained from the real data are considered to contribute significantly to classification if they have a value within the top 5% values in the 200-dimensional vector or higher. Note that this requires *every* voxel in the frequency map obtained from the real data to have a frequency value equal or higher to the top 5% *whole-brain maxima* in the frequency maps obtained from simulated data. In other words, using this threshold, in 95% of the frequency maps obtained from simulated data *zero* voxels would be above threshold. This ensures that the threshold used is corrected for multiple comparisons.

Classification of identity with generalization across stimulus modality was performed in the functionally defined ROIs using linear SVMs with flexible C parameter. The classifier was trained to distinguish between the two distractor identities with data in one stimulus modality, and was tested on the classification between the two distractor in the other stimulus modality. Accuracies were averaged and the significance of above chance classification across participants was assessed with a standard t-test.

## Results

### Responses to voices and tool sounds in face selective ROIs

A brain region encoding multimodal representations of person identity does not have to show selective responses for faces. For instance, if a region encodes representations of person identity as well as representations of other individual objects, it could be equally active during the recognition of person identity and during the recognition of other individual objects. However, several brain regions show selective responses for faces (Kanwisher et al. 1997, Sergent et al. 1992, Gauthier et al. 2000, Rajimehr et al. 2008). If a brain region encodes multimodal representations of face identity and shows selective responses for faces, we would expect it to also show selective responses for voices, because the same representations would be activated during the processing of faces and of voices. Therefore, to test whether regions showing face selectivity could encode multimodal representations of person identity, we sought to determine whether they also responded more strongly to voices that to tool sounds. Regions showing selective responses for faces were individuated with an 8-minute run in which faces and houses were presented in 16 seconds blocks while participants performed a 1-back task. The contrast between face responses and house responses yielded a network of face selective regions including the FFA bilaterally, the anterior temporal lobes bilaterally, the posterior STS (pSTS) bilaterally, and the posterior cingulate (see Table 1 for the MNI coordinates of the peaks in individual participants). In a separate 8-minute run, participants listened to voices and tool sounds, presented in 16 seconds blocks, and had to perform a 1-back task. Responses to voices and tool sounds were investigated within the face selective ROIs defined with the faces vs. houses contrast (Fig. 2A). No differences in the responses to voices and tool sounds were observed in the FFA bilaterally (right FFA t(10) = -0.36, p > 0.1; left FFA t(10) = 1.34, p > 0.1) and in the anterior temporal lobes bilaterally (right ATL t(10) = 0.85, p > 0.1; left ATL t(10) = 1.32, p > 0.1). In the right and left pSTS, responses to voices were significantly stronger than responses to tool sounds (right pSTS t(10) = 2.50, p < 0.05; left pSTS t(10) = 3.03, p < 0.05). Stronger responses to voices than tool sounds were also observed in the posterior cingulate (t(10) = 7.37, p < 0.005).

**Figure 2.**
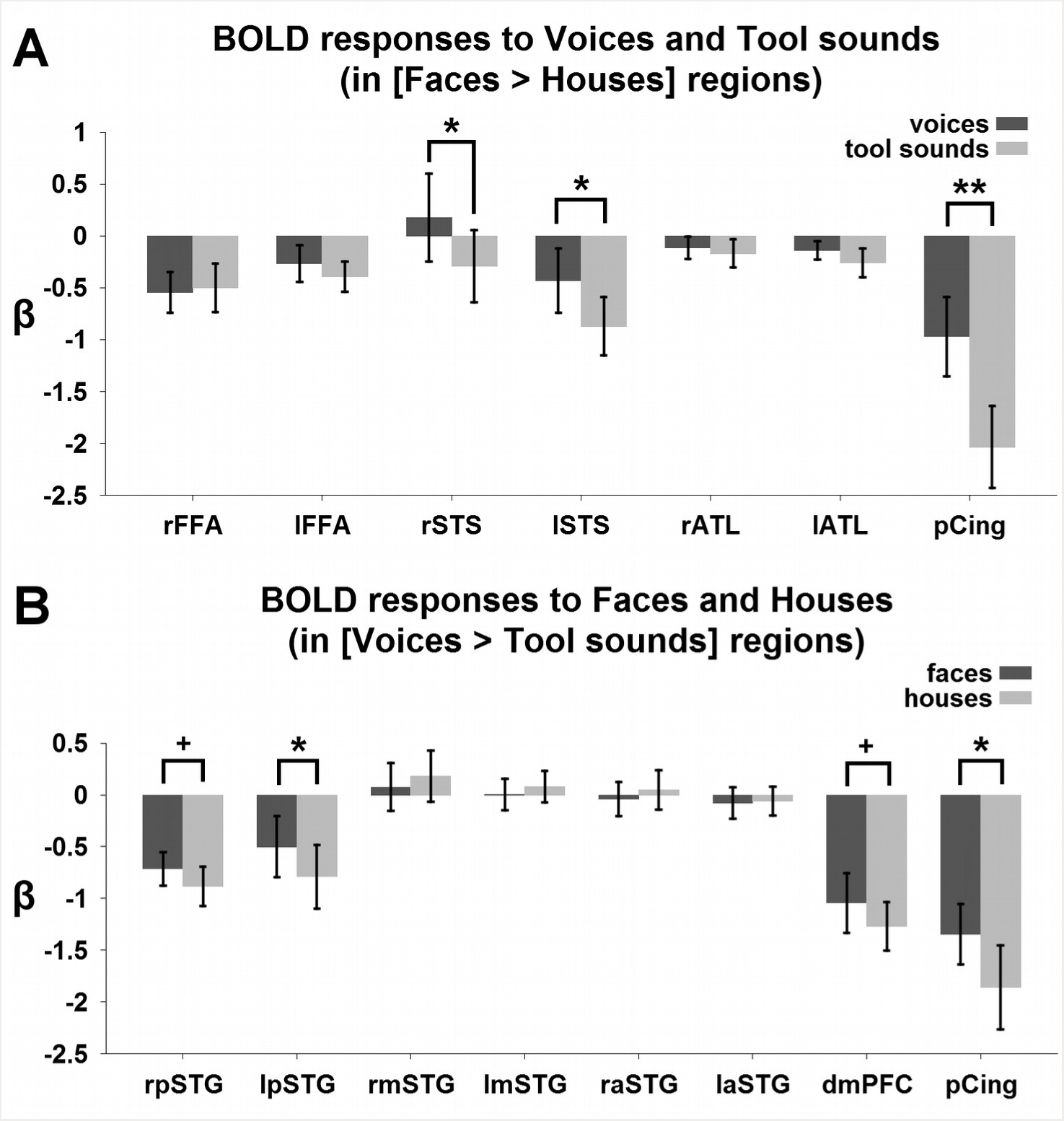
A) Mean beta values for voices and tool sounds in ROIs defined as showing stronger responses to faces than to houses. B) Mean beta values for faces and houses in ROIs defined as showing stronger responses to voices than tool sounds.

### Responses to faces and houses in voice selective ROIs

A symmetrical analysis was performed testing differential responses to faces and houses in ROIs defined with the voices vs. tool sounds contrast (Fig. 2B). The voices VS tool sounds contrast yielded 8 ROIs (see Table 2 for the MNI coordinates of the peaks in individual participants): the posterior STG bilaterally (pSTG), the mid STG bilaterally (mSTG), the anterior STG bilaterally (aSTG), ventromedial prefrontal cortex (vmPFC) and posterior cingulate. Responses to faces and houses in the aSTG and mSTG did not differ (right mSTG: t(10) = -0.69, p > 0.1; left mSTG: t(10) = -0.63, p > 0.1; right aSTG: t(10) = -0.79, p > 0.1; left aSTG: t(10) = -0.22, p > 0.1). In the left pSTG, faces elicited stronger responses than houses (t(10) = 2.64, p < 0.05), and a trend was observed in the right pSTG (t(10) = 1.96, p = 0.08). Stronger responses to faces than to houses were found in the posterior cingulate (t(10) = 2.62, p < 0.05) and a trend was found in vmPFC (t(10) = 2.21, p = 0.052).

### Recursive feature elimination mapping of face and voice information

After the two 8-minute runs consisting of blocks of faces and houses and of voices and tool sounds, participants completed 5 additional 10 minute runs in which they were presented with faces and voices and were asked to detect a target identity, regardless of the modality of the stimuli. For each of three famous Italian politicians (same nationality as the participants), we selected two face images and two audio clips of their voice (see Methods for details on the stimuli). Before entering the scanner, participants were instructed to consider one of the three identities as the target. As in previous experiments (Anzellotti et al. 2013, Anzellotti and Caramazza 2015) only the data from the distractor identities were analyzed, to avoid a confound of identity by behavioral response. In order to individuate regions that might encode multimodal representations of person identity without showing selectivity for faces or voices, two multivariate recursive feature elimination (RFE) mapping (Formisano et al. 2008, DeMartino et al. 2008) analyses were performed to individuate regions encoding information that contributes to distinguishing between face tokens depicting different people and regions encoding information that contributes to distinguishing between specific voice sounds ("voice tokens") corresponding to different people. RFE individuates voxels that contribute to classification reliably across participants and runs. The interpretation of RFE results differs to some extent from the interpretation of searchlight results because RFE guarantees that the voxels individuated contribute to the classification (while in searchlight it might be driven by neighboring voxels), but unlike searchlight in RFE a small local neighborhood of the voxel may not be itself sufficient for accurate classification (see Anzellotti and Caramazza 2014 for a more detailed discussion). The whole-brain RFE analysis in this study provides a perspective on the data which is complementary to the subsequent cross-modal classification analysis. Specifically, the RFE analysis enables to look at information encoded about faces and voices outside the functional ROIs. However, it does not test the multimodality of representations, and as other types of whole-brain analyses (e.g. searchlight) it is susceptible to between-subject variability in the anatomical location of functional regions. Replicating the results obtained in previous studies (Anzellotti et al. 2013, Anzellotti and Caramazza 2015), information that distinguishes face tokens depicting different people was found in occipital cortex, posterior temporal cortex, and the anterior temporal lobes, as well as posterior cingulate and IPS bilaterally (Fig. 3A, Fig. 4). In addition, information about face tokens depicting different people was found in the insula bilaterally (Fig. 3A, Fig. 4). This region was not individuated in previous studies using unfamiliar people as stimuli (Anzellotti et al. 2013, Anzellotti and Caramazza 2015).

**Figure 3.**
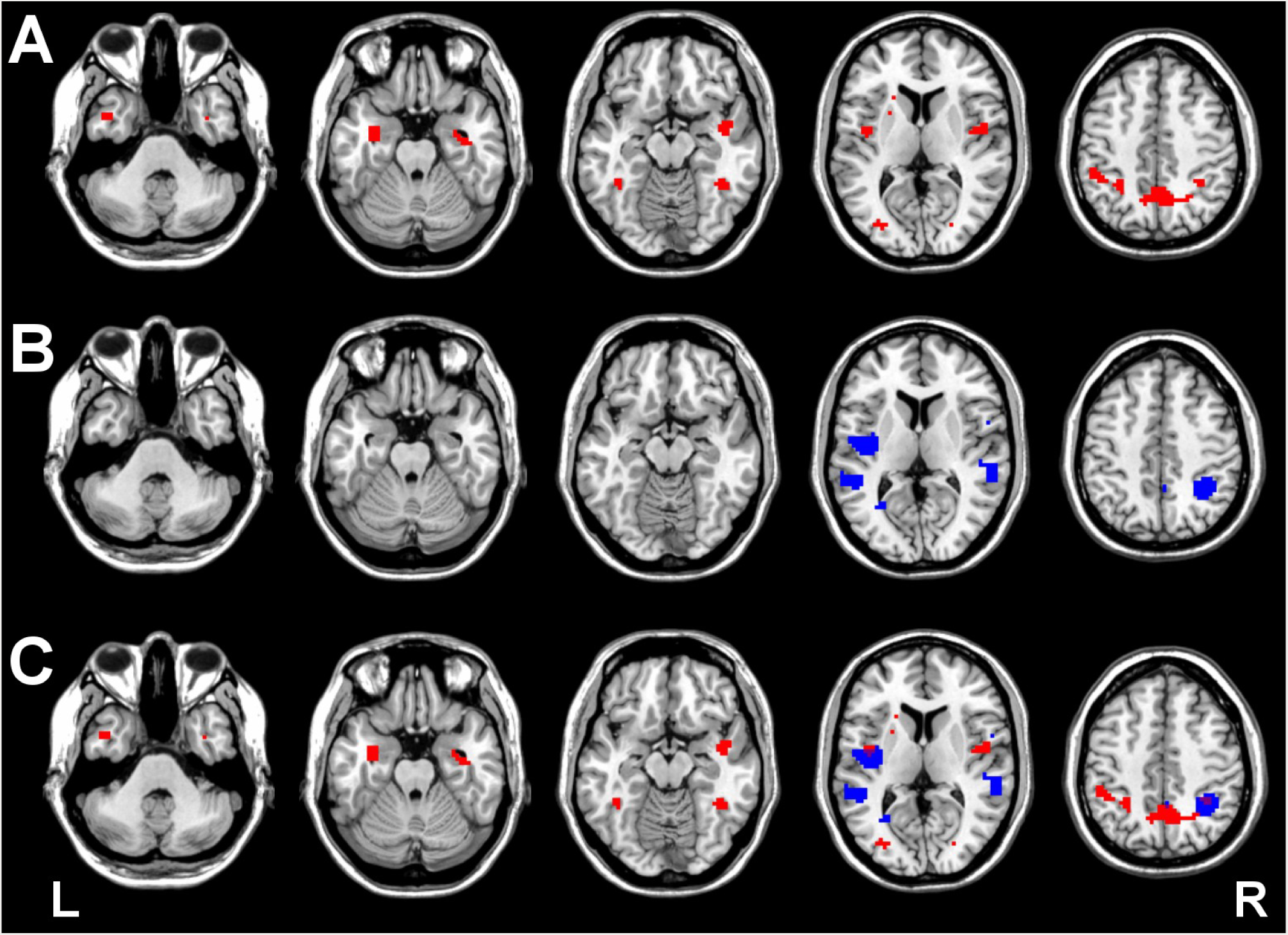
A) Regions encoding information about face tokens depicting different individuals (whole brain RFE, p < 0.05 corrected). B) Regions encoding information about voice tokens corresponding to different individuals (whole brain RFE, p < 0.05 corrected). C) Overlap between the two maps.

**Figure 4.**
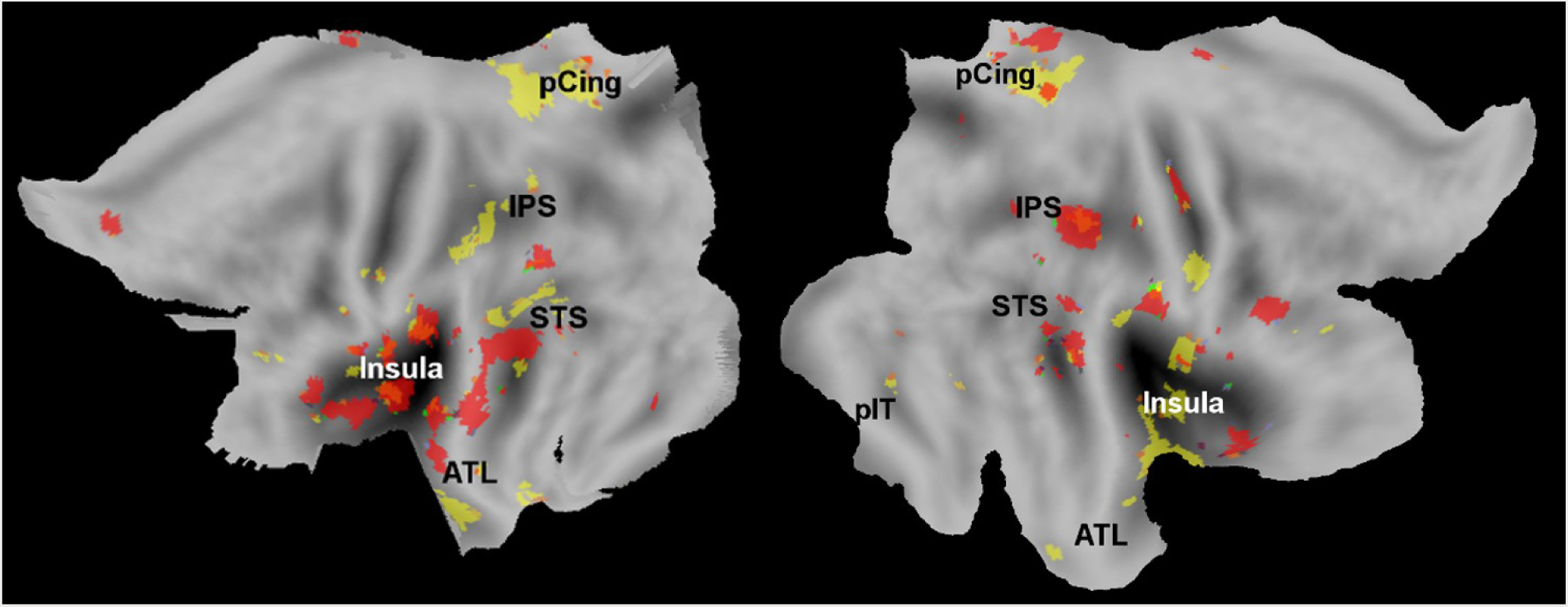
Overlap between the regions encoding information about face tokens depicting different individuals (in yellow) and the regions encoding information about voice tokens corresponding to different individuals (in red).

Information distinguishing voice tokens corresponding to different people was found in STG bilaterally, as well as posterior cingulate, right IPS, and the left insula (Fig. 3B, Fig. 4). No overlap was found in the anterior temporal lobes between regions encoding information that distinguishes between face tokens depicting different people and regions encoding information that distinguishes between voice tokens corresponding to different people (Fig. 3C, Fig. 4).

### Classification of person identity across modalities

As a strong test of the presence of multimodal representations of person identity, a classifier was trained to distinguish between the two distractor identities using the responses to stimuli in one modality and tested to distinguish the two identities with the responses to stimuli in the other modality. This analysis was performed in the ROIs defined with the faces vs. houses contrast (Fig. 5A), as well as in the ROIs defined with the voices vs. tool sounds contrast (Fig. 5B). In the face selective ROIs, classification of identity generalizing across stimulus modality was significantly above chance in the right pSTS (accuracy = 54.77%, t(10) = 4.38, p < 0.005; Bonferroni corrected p-value for 15 multiple comparisons < 0.05). This region was also found to be more active for faces than houses and for voices than tool sounds in previous analyses (Fig. 2A). However, importantly, the differences between the univariate responses to the face identities and to the voice identities in the right pSTS were non-significant (t(10) = 0.2303, p = 0.83 and t(10) = 1.97, p = 0.11) respectively), showing that the distinction between identities could not be observed in the univariate responses alone. Classification of identity generalizing across stimulus modality was not significantly above chance in the other face selective ROIs, nor was it significantly above chance in any of the voice selective ROIs (Fig. 5).

**Figure 5.**
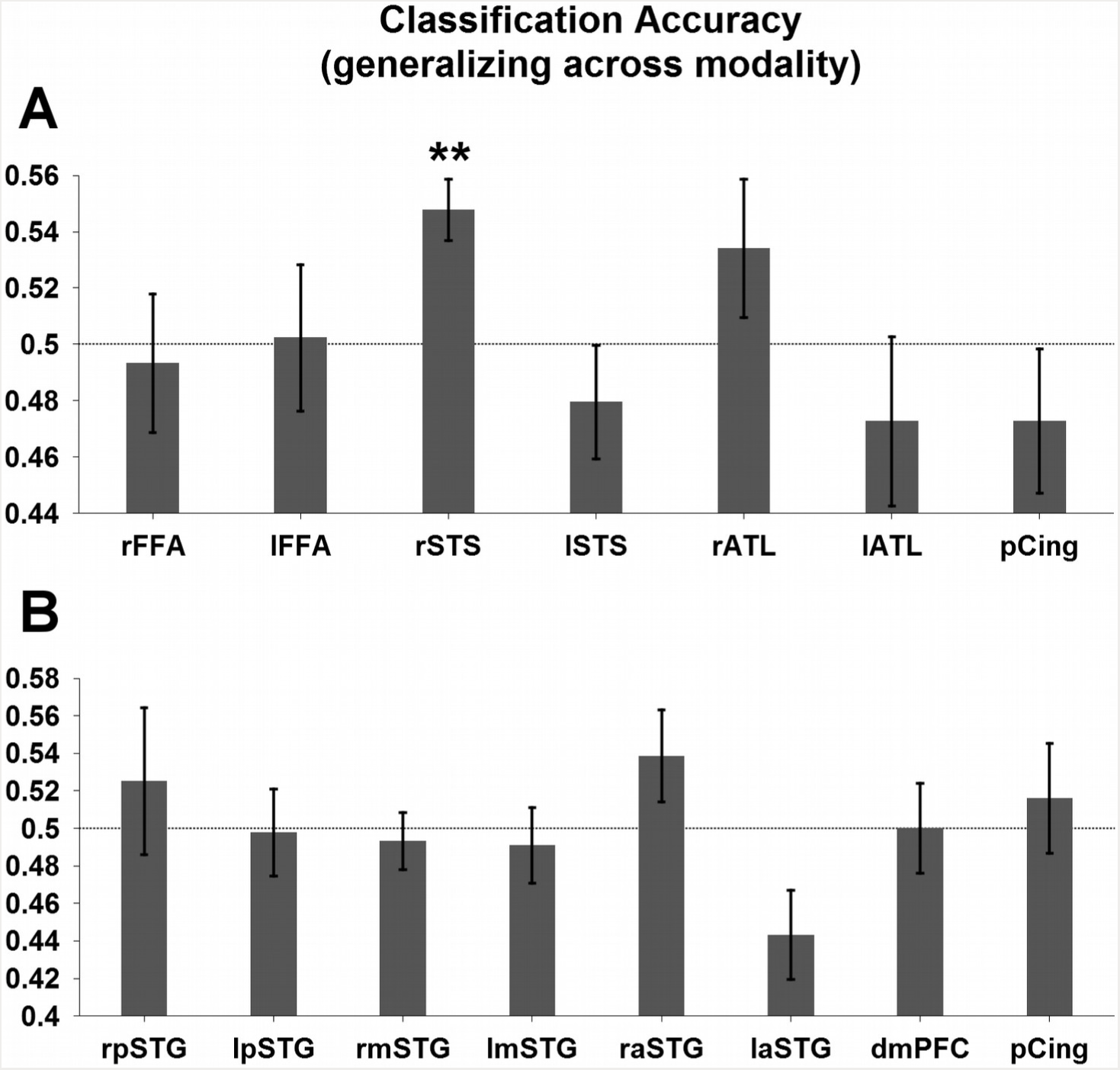
A) Classification accuracy (ratio of correct classifications) for the classification of identity training on stimuli in one modality and testing on the stimuli in the other modality in ROIs defined as showing stronger responses to faces than to houses. Significant classification was observed in the right STS (accuracy = 54.77%, t(10) = 4.38, p < 0.005). B) Classification accuracy (ratio of correct classifications) for the classification of identity training on stimuli in one modality and testing on the stimuli in the other modality in ROIs defined as showing stronger responses to voices than tool sounds. None of the voice selective ROIs was found to achieve significant cross-modal classification of identity.

### Classification of person identity within modality in the pSTS

If pSTS encodes representations of person identity, it should also classify person identity generalizing across transformations within the visual modality. To test this hypothesis, we reanalyzed the data from a previous experiment in which we investigated classification of person identity generalizing across viewpoints (Anzellotti et al. 2013), but in which right pSTS had not been included as a region of interest. Since pSTS could not be localized in each individual participant in that study, we defined pSTS as a spherical ROI centered in the pSTS group peak for the contrast faces vs houses. We trained a support vector machine to classify person identity training with 4 out of 5 viewpoints and tested the classification accuracy on the fifth viewpoint, this procedure was iterated for all choices of the left-out viewpoint and the accuracies were averaged. Classification of person identity generalizing across viewpoint in the right pSTS was significant (accuracy = 54.91%; t(8) = 2.56, p = 0.034).

### Summary of results

Anterior temporal lobe face selective regions do not show selective responses for voice stimuli, and anterior temporal lobe voice selective regions (aSTG) do not show selective responses for face stimuli. Furthermore, no overlap between information maps for face tokens and voice tokens was found in the anterior temporal lobes. Finally, classification of person identity generalizing across stimulus modality in the anterior temporal lobe ROIs was at chance. Evidence of selectivity for both faces and voices was found in posterior cingulate and in pSTS bilaterally. The responses in right pSTS allowed classification of person identity generalizing both across transformations within the visual modality, and across different stimulus modalities.

## Discussion

In the present study, we found that the face-selective right pSTS shows voice-selectivity, and encodes information about person identity generalizing across different stimuli within and across modality. The pSTS is known to integrate visual and auditory information both in humans (Grossman et al. 2000) and in monkeys (Vaina et al. 2001; Grossman et al. 2005). However, the pSTS is often implicated in the representation of biological motion (Haxby et al. 2000; Calder and Young 2005; Narumoto et al. 2001).

In the literature on face perception it has been proposed that ventral temporal regions are mostly involved in processing static aspects of faces such as identity, while the pSTS is mostly involved in processing dynamic aspects such as facial expressions (Winston et al. 2004). Why, then, did we find identity information in the pSTS? The view that pSTS is exclusively involved in processing dynamic aspects of faces is not strongly supported by the empirical evidence (Calder and Young 2005). Narumoto et al. (2001) found stronger responses in the right pSTS when participants attended to the expression of faces, but this does not rule out an involvement of pSTS in the representation of identity. In line with the view that pSTS represents information about person identity, Winston et al. (2004) found fMR adaptation (fMR-A) in pSTS for repetitions of face identity. Adaptation for repetitions of the same face identity in the pSTS was even stronger than adaptation for repetitions of the same facial expression (Winston et al. 2004). Stronger effects for repetition of facial expression than for repetition of face identity occurred only in a more anterior STS region (Winston et al. 2004).

The right pSTS has been previously implicated in the processing of voices (Belin, Zatorre, & Ahad, 2002; Belin et al. 2000), and hypothesized to play a role in face-voice integration because of its functional connectivity with FFA (von Kriegstein et al. 2005). Furthermore, electrophysiology research in macaques has shown that simultaneous presentation of faces and voices leads to interaction effects between auditory cortex and STS which are not observed within auditory cortex (Ghazanfar, Chandrasekaran & Logothetis 2008). This effect is not general to all kinds of visul stimuli, as it was not observed for simultaneous presentation of a voice and a disk (Ghazanfar, Chandrasekaran & Logothetis 2008).

A parsimonious account of the present results holds that pSTS encodes multimodal representations. However, it could be questioned whether voice selectivity and classification of identity across modalities could arise in pSTS via other mechanisms, such as the reactivation of face representations triggered by the voice, especially given tractography evidence pointing to direct connections between regions responding to faces and voices (Blank, Anwander & von Kriegstein 2011). A hypothesis proposing generic reactivation of face representations in response to familiar voices does not account well for the present data, because neither voice-selective responses nor classification of identity across modality were observed in face-selective regions other than the pSTS (FFA, ATL). Despite this, the possibility that face representations restricted specifically to the pSTS were reactivated during voice recognition is difficult to rule out.

An important question concerns the afferents of pSTS: where does the visual and auditory information represented in pSTS come from? Studies using retrograde tracers in the posterior STS of rhesus monkeys labeled the anterior temporal lobe, the lower bank of IPS, and the posterior cyngulate gyrus (Seltzer and Pandya 1994, see cases 12, 14). This network includes regions of overlap between the RFE maps for faces and voices in our study (IPS and posterior cingulate) as well as the anterior temporal lobes. Visual and auditory information could be conveyed from the anterior temporal lobes to pSTS, while afferents from IPS could contribute to the allocation of attentional resources (Xu and Chun 2009).

Unlike in the pSTS, no evidence of classification of identity across modality was found in the ATL. However, caution should be exerted in interpreting null results, especially in analyses requiring fine-grained classification. Multimodal representations of person identity might exist in ATL but the spatial distribution of neurons encoding different identities might prevent decoding with MVPA (see Dubois, de Berker and Tsao 2015 for a discussion of this possibility). The univariate analyses in this study show that separate regions within ATL show selectivity for faces and voices, indicating that within ATL object representations are organized by stimulus modality. The more ventral location of responses to visual stimuli and the more dorsal location of responses to auditory stimuli is consistent with a meta analysis of neuroimaging studies reporting ATL activity (Visser et al. 2010).

Studies of patients with semantic dementia (SD) have provided a wealth of evidence on the consequences of damage to the anterior temporal lobes (Snowden et al. 1989; Hodges et al. 1992; Thompson et al. 2003). Semantic dementia patients with deficits for face recognition usually also have deficits for voice recognition (Snowden et al. 1989; Gainotti et al. 2004). By contrast, recognition of people from their names and recognition from their faces are often dissociated, with patients affected by left temporal variant SD showing greater impairment for recognition of people from their names, and patients affected by right temporal variant SD showing greater impairment for recognition of people from their faces (Snowden et al. 2004).

These results seem to suggest that separate regions are involved in representing people's names and people's faces and voices, but that the same regions are involved in representing both faces and voices. However, in neuropsychology associations are less informative than dissociations (Caramazza 1984): an association could result from simultaneous damage to two different structures, in this case ventral ATL and the anterior STG. Consistent with this account, voxel based morphometry research (Gorno-Tempini et al. 2004) found reduced cortical gray matter in both ventral portions of the right temporal pole and the right anterior STG in a patient affected by right temporal variant SD. An analogous pattern was observed in studies of left temporal variant SD, with decreases in cortical gray matter volume in both ventral anterior temporal regions and the anterior STG (Mummery et al. 2000). More generally, SD seems to damage ventral and dorsal portions of the anterior temporal lobes to similar extents, while it damages the left and right hemispheres to different extents (Mummery et al. 2000; Thompson et al. 2003; Snowden et al. 2004). As a consequence, cognitive abilities implemented by regions that are differentiated by hemisphere would tend to show dissociations, while cognitive abilities implemented by regions that are differentiated along the ventral-dorsal axis would tend to show associations. Therefore, the neuropsychological evidence of joint face and voice impairment in SD patients is compatible with the possibility of separate representations of faces and voices in ATL. In further accord with this possibility, recent research reported dissociations between face recognition and voice recognition deficits in patients with ATL damage (Liu et al. 2014).

The RFE maps for information about individual faces and about individual voices were found to overlap in the IPS, suggesting that also IPS could encode information about identity generalizing across modalities. IPS has been previously implicated in object identification and tracking (Xu & Chun, 2006, 2007, 2009; Xu, 2007, 2008, 2009, Jeong and Xu 2013), and it could play a similar function for person identity. Since IPS could not be localized with the independent localizers in this study, a ROI analysis could not be performed for IPS. Another region showing overlap between the RFE maps for information about individual faces and about individual voices was the left insula. We speculate that responses in this region may depend on emotional reactions to the persons that are common to both modalities (Phan et al. 2004, Lamm and Singer 2010). However, additional studies are needed to clarify the roles of IPS and the insula during the recognition of person identity.

The present study found that pSTS encodes representations of person identity that generalize across stimulus modality. Evidence of organization by modality of person representations was found in ATL, with face representations in ventral regions and voice representations in more dorsal regions. We speculate that person identity information reaches pSTS via afferent fibers originating in the ATL.

## Acknowledgements

We thank Valentina Brentari for assistance in collecting data, Hochan Lee and Alan Rozet for assistance in the preparation of the stimuli. This work was supported by the Provincia Autonoma di Trento and by the Fondazione Cassa di Risparmio di Trento e Rovereto. S. A. was supported by a Dissertation Completion Fellowship from Harvard University and from a Fellowship from the Simons Center for the Social Brain.

